# Unveiling the double-edged sword: SOD1 trimers possess tissue-selective toxicity and bind septin-7 in motor neuron-like cells

**DOI:** 10.1101/2024.03.22.586186

**Authors:** Esther Sue Choi, Brianna Leigh Hnath, Congzhou Mike Sha, Nikolay V Dokholyan

**Affiliations:** Department of Pharmacology, Penn State College of Medicine, Hershey, PA, USA; Medical Scientist Training Program, Penn State College of Medicine, Hershey, PA, USA; Department of Biomedical Engineering, Penn State University, University Park, PA, USA; Department of Biochemistry and Molecular Biology, Penn State College of Medicine, Hershey, PA, USA; Department of Chemistry, Penn State University, University Park, PA, USA

**Author notes:** HCAR 3049 1214 Research Blvd Hummelstown, PA 17036.

**Keywords:** SOD1 oligomers, SOD1 trimer, protein-protein interactions, interactome, septin-7, neurodegeneration

## Abstract

Misfolded soluble trimeric species of superoxide dismutase 1 (SOD1) are associated with increased death in neuron-like cell models and greater disease severity in amyotrophic lateral sclerosis (ALS) patients compared to insoluble protein aggregates. The mechanism by which structurally independent SOD1 trimers cause cellular toxicity is unknown but may be a driver of disease pathology. Here, we uncovered the SOD1 trimer interactome – a map of potential tissue-selective protein binding partners in the brain, spinal cord, and skeletal muscle. We identified binding partners and key pathways associated with SOD1 trimers, comparing them to those of wild-type SOD1 dimers. We found that trimers may affect normal cellular functions such as dendritic spine morphogenesis and synaptic function in the central nervous system and cellular metabolism in skeletal muscle. We also identified key pathways using transcriptomic data from motor neuron-like cells (NSC-34s) expressing SOD1 trimers. We discovered differential gene expression in cells that express SOD1 trimers with selective enrichment of genes responsible for protein localization to membranes and a global upregulation of cellular senescence pathways. We performed detailed computational and biochemical characterization of protein binding for septin-7, an SOD1 trimer binding partner. We found that septin-7 preferentially binds SOD1 trimers and co-localizes in neuron-like cells. We explore a double-edged sword theory regarding the toxicity of SOD1 trimers. These trimers are implicated in causing dysfunction not only in the central nervous system but also in muscle tissues. Our investigation highlights key protein factors and pathways within each system, revealing a plausible intersection of genetic and pathophysiological mechanisms in ALS through interactions involving SOD1 trimers.

**Summary:** In amyotrophic lateral sclerosis (ALS), misfolded soluble species of superoxide dismutase 1 (SOD1) are associated with disease severity and, specifically, trimeric forms of SOD1 are toxic in neuron-like cells compared to insoluble aggregates. The role of toxic SOD1 trimers in cells is unknown. Using molecular engineering and pull-down experiments, we found that SOD1 trimers have tissue-selective protein interactions that affect pathways such as dendritic spine morphogenesis and synaptic function in the nerves, energy, and amino acid metabolism in skeletal muscle. We investigated the SOD1 trimer transcriptome to reveal a global upregulation of genes associated with cellular senescence compared to SOD1 dimers. We further validated septin-7, a shared brain and spinal cord protein binding hit, using integrative computational and biochemical approaches, and confirmed that septin-7 binds SOD1 trimers and not native dimers. Taken together, we show evidence that SOD1 trimers play a central role in the convergence of ALS pathophysiology.

**Graphical abstract:** 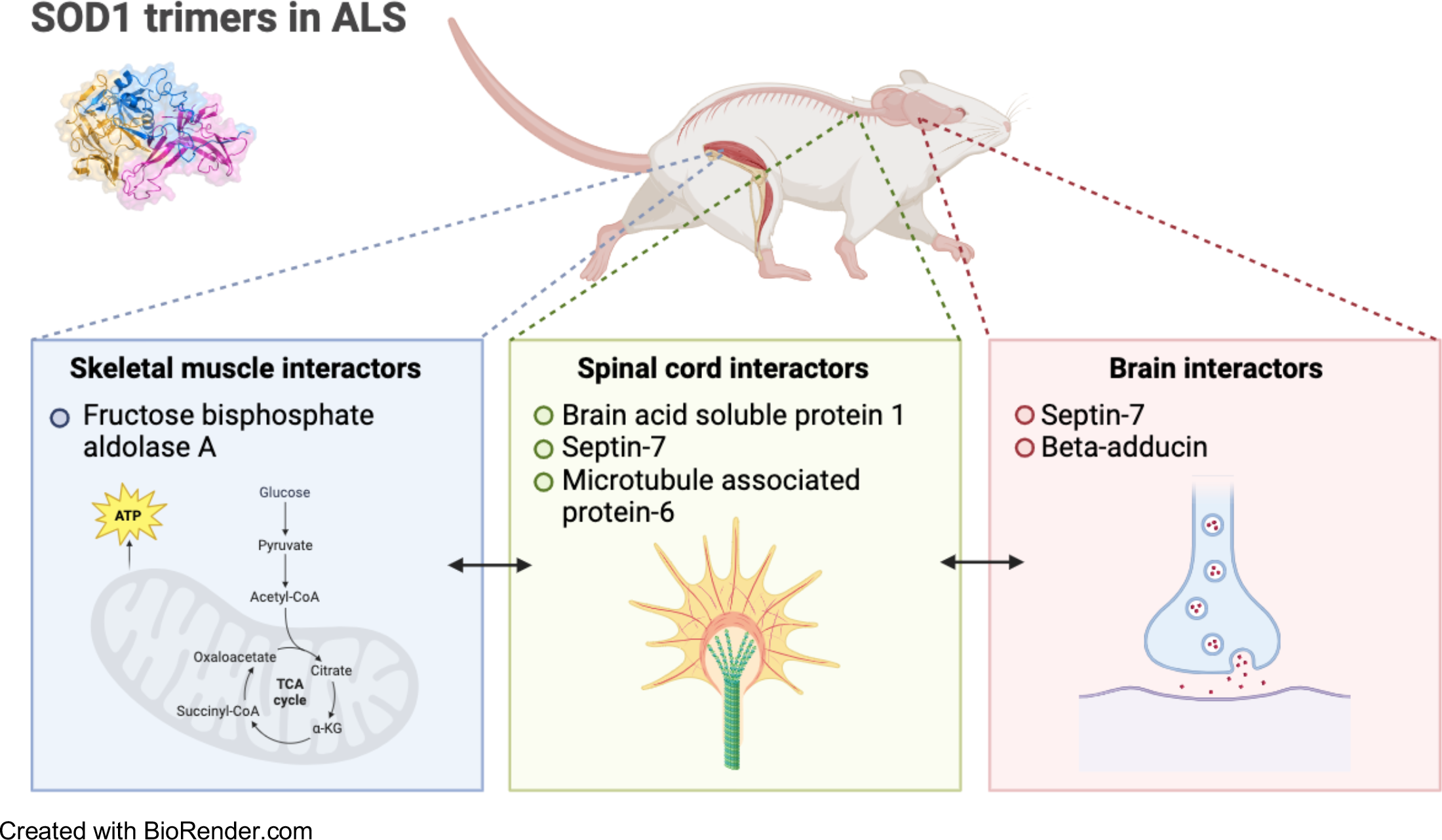

## Introduction

Amyotrophic lateral sclerosis (ALS) is a neurodegenerative disease that targets both upper and lower motor neurons in the central nervous system (CNS) leading to progressive muscle weakness and atrophy^1^. The incidence of ALS is increasing due to aging populations, and the disease remains fatal, with a mean survival after diagnosis of 2-4 years^2^. Over the past decades, over 50 potential causative or disease-modifying genes have been identified for ALS^3^. About 10% of ALS diagnoses have a known genetic cause and mutations in superoxide dismutase 1 (SOD1) comprise of about 20% of familial ALS. For the protein SOD1, most mutations are missense mutations, resulting in a wide range of phenotype severity^4^. In addition, SOD1 may be involved in ALS pathogenesis even for patients lacking mutations in SOD1^5–8^. Whether genetic factors such as SOD1 mutations play a role in sporadic ALS is still highly debated and the exact mechanism in familial cases is unknown^9,10^. Regardless of the origin, Cleveland and Rothstein^11^ presented four principals that may account for neuron death: oxidative damage, axonal strangulation from disorganized neurofilaments, toxicity from intracellular aggregates, and excitotoxic death from glutamate mishandling. SOD1 is known to have a gain-of-function^12,13^ mechanism in ALS, which is why it is crucial to investigate its direct protein interactions. Some mechanisms proposed for SOD1 in ALS include excitotoxicity^14^, oxidative stress^15^, endoplasmic reticulum stress^16,17^, and mitochondrial dysfunction^18^. It is unknown how SOD1 mutations cause clinical symptoms, but the variety of proposed mechanisms for SOD1-related ALS illuminates the complexity of ALS pathophysiology. Uncovering the molecular mechanism of SOD1 is not limited to just SOD1-related ALS. Despite extensive research efforts, much about the precise triggers and progression of ALS remains elusive.

Mounting evidence suggests that small, soluble misfolded oligomers are the driver of disease^19^. Evidence for SOD1 oligomer toxicity exists on molecular, cellular, and organismal levels^20,21^. Proctor and colleagues^22^ showed that a small soluble aggregate of SOD1, the trimer, is toxic in an NSC-34 cell model of ALS. Through experimental and computational methods, Proctor et al.^22^, generated 3D molecular models of SOD1 oligomers to establish stoichiometry, structural components, and critical residues for SOD1 trimer stability. The proposed SOD1 trimer model was validated in a neuronal cell line showing an association of increased cell death with SOD1 trimer stability compared to wild-type SOD1. Destabilizing the SOD1 trimer showed decreased cell death, further confirming the toxicity of SOD1 trimers compared to the native dimer. Zhu et al.^23^ further compared the cytotoxicity of SOD1 trimers with that of larger non-soluble SOD1 aggregates; large aggregates did not impact cell death, whereas trimers were still highly associated with cell death. They designed SOD1 mutants that stabilized either the larger aggregate form, SOD1 fibrils, or SOD1 trimers. Trimer-stabilizing mutants induced increased neuronal death compared to fibril-stabilizing mutants, which instead attenuated cell death. Additionally, in mice models of ALS, Gill et al.^24^ found that levels of SOD1 aggregation in spinal cords were inversely correlated with disease progression. Areas that were highly affected by disease contained high levels of soluble, misfolded SOD1. Recently, Hnath et al.^25^, discovered that toxic SOD1 trimers form off-pathway and directly compete with the formation of larger oligomers and protective fibril formation. SOD1 trimers are structurally distinct from larger oligomers and aggregates suggesting that SOD1 trimers may have independent binding partners and it may be crucial to parse out selective protein interactions of toxic SOD1 trimers. Overall, there is strong evidence that SOD1 trimers are structurally different than other forms and the molecular species responsible for cell death in SOD1- related ALS while larger aggregates and fibrils are not harmful^4,26–28^. The paradigm in neurodegeneration has shifted from attributing toxicity to aggregates to recognizing soluble oligomers as the toxic entities, given their association with increased disease severity and death^25^. Attributing neurotoxicity to a trimeric protein species versus aggregates revolutionizes our understanding of neurodegeneration, where previous efforts, such as in Alzheimer’s disease and Parkinson’s disease, are focused on preventing the formation of and disrupting protein aggregates^29^.

The molecular mechanism of SOD1 trimer toxicity is unknown, hence, we focus on proteins that directly interact with the trimers to reveal downstream pathways leading to cytotoxicity. We hypothesize that toxic SOD1 trimers bind tissue-selective proteins to induce cell dysfunction ultimately leading to death of motor neurons. To this end, we performed trimer-binding pulldown assays, mass spectrometry proteomics, and RNA transcriptome analysis to identify potential SOD1 trimer binding partners and critical cell pathways associated with trimers. We then confirmed the interaction of the SOD1 trimer with binding partner septin-7 using molecular dynamics simulations, microscale thermophoresis, and confocal microscopy with colocalization analysis. Understanding the binding partners of SOD1 trimers allow us to shed light on the downstream pathways and we propose that SOD1 and septin-7, previously genetically linked to ALS, may belong to the same pathological pathways leading to motor neuron death.

## Results

### SOD1 trimers bind tissue-selective proteins in the brain, spinal cord, and skeletal muscle

We identified potential protein binders of SOD1 trimer in wild-type (WT) mice brain, spinal cord, and skeletal muscle. We performed pull-down experiments by crosslinking purified native SOD1 dimers or mutant SOD1 trimer proteins to magnetic carboxyl beads and then exposed the SOD1-beads to the mouse tissue lysates. We pulled down the proteins that bound to either SOD1 dimers or SOD1 trimers and identified the proteins with mass spectrometry (Figure 1a). Then, we built protein-protein interaction (PPI) maps using STRING.db^30,31^. SOD1 trimers pulled down selective proteins of various sizes from the three different tissue types visualized by silver-stained SDS-PAGE gel compared to WT SOD1 dimers (Figure 1b). We observed a difference in proteins bound by trimers compared to dimers based on tissue type. Our mass spectrometry analysis comparing protein interactors of SOD1 timers and WT SOD1 dimers resulted in 283 total protein hits in the brain, 63 total protein hits in skeletal muscle, and 33 total protein hits in the spinal cord (Figure 1c). Of the total brain protein hits, 129 had a positive fold change indicating that these proteins bound SOD1 trimer greater than they bound SOD1 dimer. For the skeletal muscle there were 17 positive fold change hits and in the spinal cord there were 25 proteins. We performed Mann-Whitney statistical testing with adjusted p-value cutoffs (at the false positive rate *α* = 0.05 level of significance) within each tissue using the Benjamini-Hochberg procedure, and found that SOD1 trimers selectively and significantly bound one protein in the brain (septin-7; p = 0.00045), five proteins in the spinal cord (cytochrome c oxidase; p = 0.0012, creatine kinase; p = 0.00012, brain acid soluble protein 1; p = 0.0019, microtubule-associated protein 1; p = 0.018, and hemoglobin sub unit alpha; p = 0.018) and one protein in skeletal muscle (fructose-bisphosphate aldolase A; p-adjusted <0.0001) (Table 1). Notably, septin-7^32–34^, brain acid soluble protein 1^35,36^, and fructose-bisphosphate aldolase A^37,38^ are tissue-specific protein hits and have been previously linked to neurodegenerative processes. Overall, SOD1 trimers feature tissue-selective interactomes compared to that of native SOD1 dimers.

**Figure 1.**
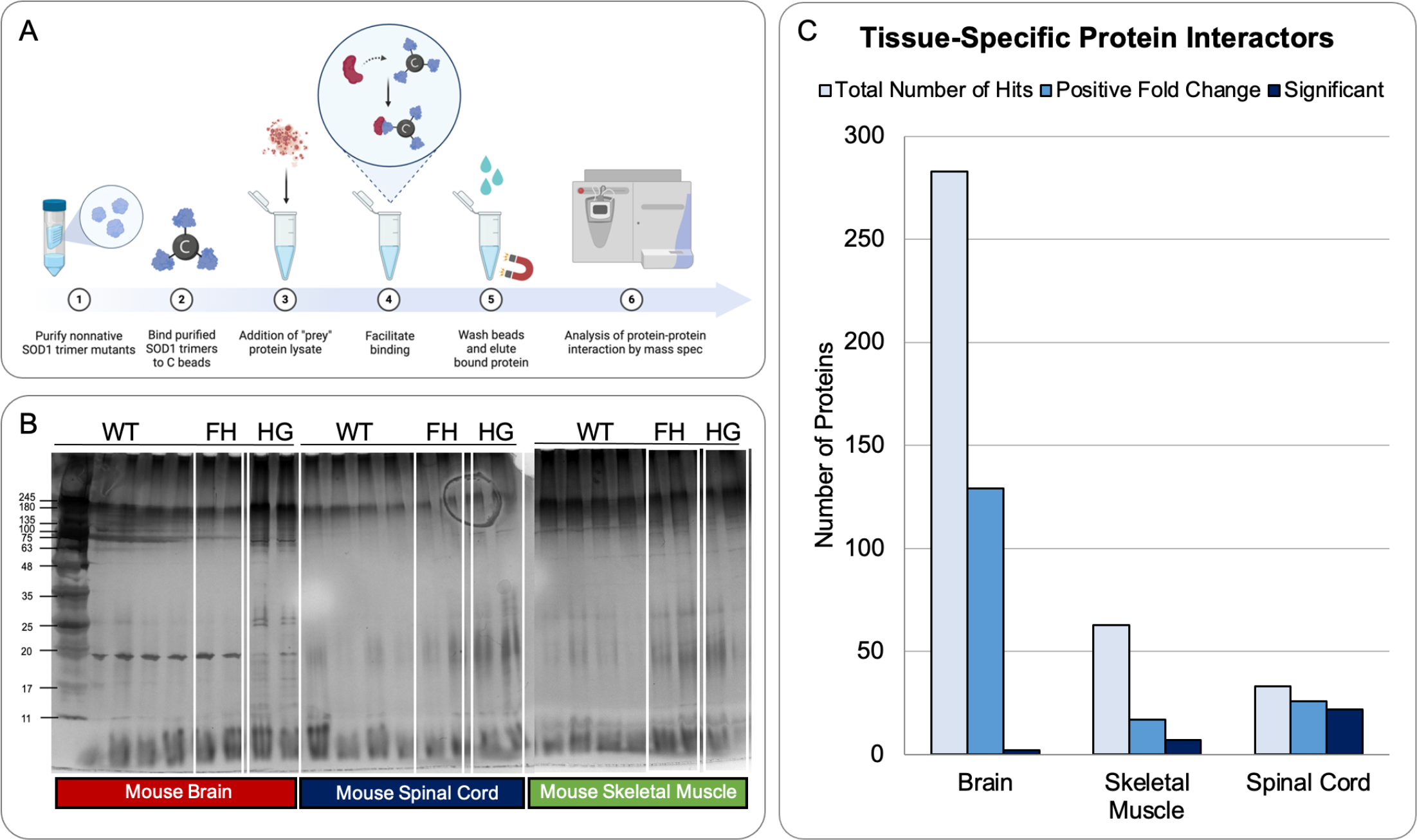
SOD1 pulldown assay overview. A) Methodology for SOD1 pulldowns with purified WT mouse tissues (created with BioRender.com). We purified SOD1 trimers and covalently bonded the trimers to magnetic carboxyl beads before exposure to mouse brain, spinal cord, or muscle tissue. Then, we performed wash steps and eluded trimer binding proteins for mass spectrometry analysis. B) Silver stained SDS-PAGE gel of pulldown elutions showing differences in protein content from mouse brain, spinal cord, and skeletal muscle. C) Bar graph displaying number of total proteins for each tissue type as well as the number of proteins that had a positive fold change and were statistically significant compared to native WT dimers.

**Table 1.** S0D1 trimer interactor proteins stratified by tissue type.

| **Table 1. SOD1 trimer interactor proteins stratified by tissue type** |  |  |  |
| --- | --- | --- | --- |
| **Tissue** | **Protein** | **Mann-Whitney test (p-value)** | **Log Fold Change (mutant/WT)** |
| Brain | Septin 7 | 0.00045 | 2.12 |
| Spinal cord | Cytochrome c oxidase | 0.00012 | 7.29 |
| Spinal cord | Creatine kinase | 0.00055 | 2.60 |
| Spinal cord | Brain acid soluble protein 1 | 0.0019 | 58.13 |
| Spinal cord | Microtubule-associated protein 1 | 0.018 | 8.07 |
| Spinal cord | Hemoglobin subunit alpha | 0.019 | 1.16 |
| Skeletal muscle | Fructose-bisphosphate aldolase A | <0.0001 | 2.14 |

### SOD1 trimer brain interactome centers around proteins associated with microtubule function, axonal transport, cytoskeletal dynamics, and synapse maintenance

Motivated by the intricate and dynamic landscape of protein interactions associated with ALS, we began by mapping the proteins identified from SOD1 trimer pulldowns with brain tissue lysate using STRING.db (Figure 2a). As we explored the interactome, distinct major groups emerged, each bearing responsibility for crucial cellular functions. The major groups are the ribosomal protein (RPL) family pivotal for protein folding, stood out alongside the ubiquinol-cytochrome (UQCRQ) group contributing to mitochondrial function Simultaneously, proteins linked to tubulin and cytoskeletal maintenance played a significant role in the observed clusters. Amidst this diversity of protein families and biological processes, microtubule-associated protein tau (MAPT) was centered in the network, emphasizing its integral position within the interactome (Figure 2a). We plotted the brain interactome hits on a volcano plot to visualize the fold change and significance of the hits binding to the SOD1 trimer versus the native dimer (Figure 2b). The red dots show proteins with a positive fold change indicating a greater binding preference to trimers. Septin-7 was statistically significant in the binding preference to SOD1 trimers compared to WT dimers with a positive fold change of 2.12 (p-value 0.00045). Beta-adducin (Add2) featured the largest fold change of 16.4 among brain hits. Of the total 283 brain hits, we find the 129 positive fold change hits (Figure 2c).

**Figure 2.**
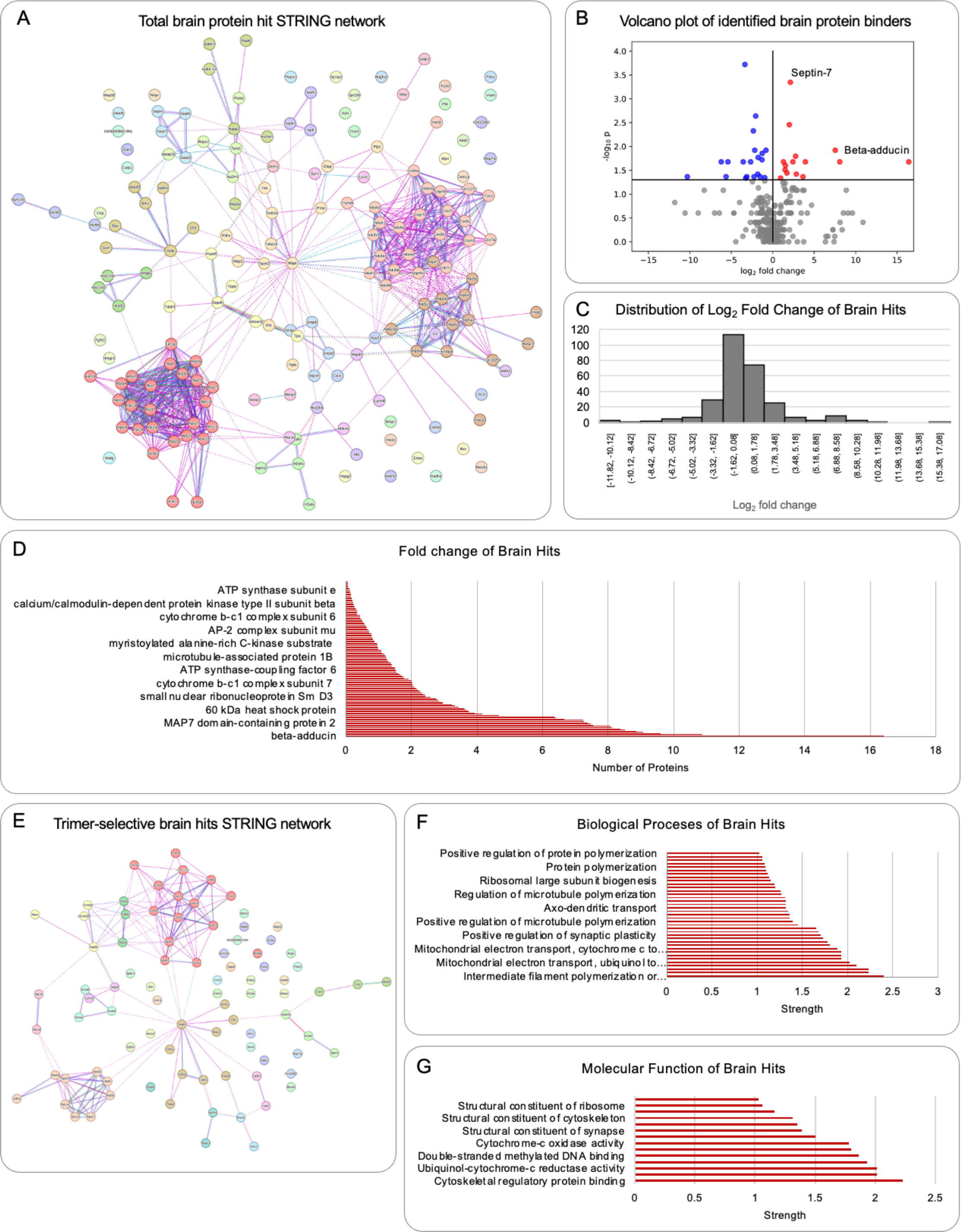
SOD1 trimer brain interactome. A) Protein-protein interaction network of SOD1 trimer hits with WT mouse brain tissue. B). Volcano plot of SOD1 trimer brain hits. Blue dots are positive fold change nodes and red dots are negative fold change nodes. Septin-7 was the most statistically significant hit with beta-adducin the hit with the greatest fold change. C). Distribution of fold change of brain hits showing normal distribution. D) Fold change of all the brain hits ordered from least change to the most change at the bottom. E) Protein-protein network of positive fold change hits displaying similar clusters to the total network map in A). F) Biological processes of the brain hits ordered by strength. G) Molecular function of brain hits ordered by strength.

To further discern the distinctions between proteins binding to SOD1 trimers and dimers, we pruned out proteins without a positive fold change from the total interactome (Figure 2d). The resultant analysis revealed that major protein families remained, with a central focus on MAPT (Figure 2e). The clusters that remained are the RPL protein family, the tubulin protein family, and the COX/UQCR protein family. Our investigation extended beyond protein clusters to encompass associated biological processes within the interactome. Processes such as intermediate filament polymerization or depolymerization, neurofilament bundle assembly, mitochondrial electron transport, and regulation of tubulin deacetylation came to the forefront (Figure 2f). The molecular function of cytoskeletal regulatory protein binding emerged predominantly within the interactome, as evidenced by its substantial strength measurement of 2.23 (Figure 2g). In interpreting these findings, the strength measurement, based on Log10 (observed/expected), offered a quantitative perspective on the enrichment effect within the interactome. This ratio provided insights into the significance of the observed proteins in comparison to what would be expected in a random network of similar size. In essence, our work has unraveled a comprehensive view of the SOD1 trimer brain interactome, emphasizing its associations with microtubule function, axonal transport, cytoskeletal dynamics, and synapse maintenance. Exploration of brain-specific protein binders contributes valuable insights to the broader understanding of cellular processes in ALS and allows for investigations into the functional implications of these protein interactions within the tissue type.

### SOD1 trimer spinal cord interactome contains proteins associated with peptidyl-cysteine S-trans- nitrosylation, microtubule, and cytoskeleton organization

We explored the SOD1 trimer protein binders in the spinal cord tissue lysate to distinguish protein interactions within the spinal cord and to shed light on both shared and unique pathways compared to the brain hits. From WT mice spinal cord lysate, we identified 27 binding proteins that underpin crucial cellular functions (Figure 3a). There are four larger clusters within the spinal cord interactome: tubulin proteins, metabolic proteins (glyceraldehyde-3-phosphate dehydrogenase, triosephosphate isomerase 1, enolase 1, creatine kinase m-type, S100 calcium-binding protein A9), ATP synthase proteins, and cytochrome c oxidase proteins (Figure 2b). The presence of septin-7 and MAPT proteins further enriched the complexity of these interactions. We observed that myelin basic protein had a fold change of 0.98 (p = 0.0001), cytochrome c oxidase had a fold change of 7.29 (p-value 0.00012), and creatine kinase had the most significant difference binding to SOD1 trimers compared to WT SOD1 dimers with a fold change of 7.29 (p-value 0.00055), but brain acid soluble protein 1 (fold change, 58.13; p = 0.0019) and septin-7 (fold change 41.73; p = 0.039) had the largest fold change in binding. The skewed distribution of log fold changes for the spinal cord hits, likely influenced by the limited number of total hits (33) revealed both subtle variations and distinct outliers (Figure 3d). We observed two with very large fold changes mentioned earlier (BASP1 and SEPT7). Subsequently, we mapped the protein-protein interactors with a positive log fold change in trimers compared to dimers and there were 3 main clusters: mitochondrial, metabolic, and structural (Figure 3e). Within this protein-protein interaction (PPI) network, GAPDH emerged as a central player. We observed a large gap between the top 2 proteins (BASP1 and septin-7) and the rest of the interactors in the log fold change of spinal cord hits.

**Figure 3.**
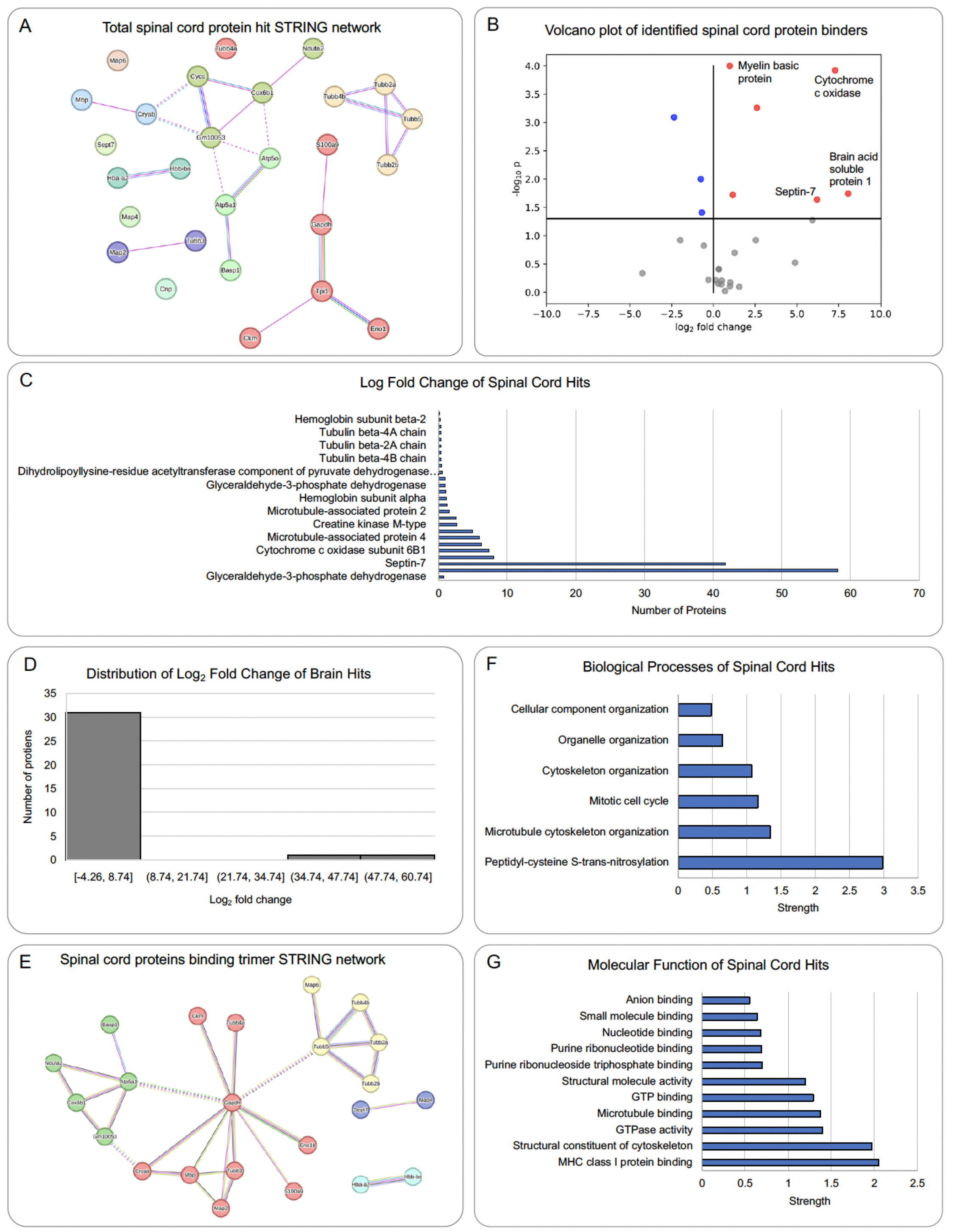
SOD1 trimer spinal cord interactome. A) Protein-protein interaction network of SOD1 trimer hits with WT mouse spinal cord tissue. B). Volcano plot of SOD1 trimer spinal cord hits. Blue are positive fold change nodes and red are negative fold change nodes. Myelin basic protein and cytochrome c oxidase were the most statistically significant hit with brain acid soluble protein 1 and septin-7 the hits with the greatest fold change. C). Distribution of fold change of spinal cord hits. D) Fold change of all the spinal hits ordered from least change to the most change at the bottom. E) Protein-protein network of positive fold change hits displaying three major cluster groups centralized by glyceraldehyde-3- phosphate dehydrogenase. F) Biological processes of the spinal cord hits ordered by strength. G) Molecular function of spinal cord hits ordered by strength.

We mapped the identified interacting partners of SOD1 trimers to biological processes, our analysis revealed peptidyl-cysteine s-nitrosylation as the top process, demonstrating a substantial strength of 2.98. Microtubule cytoskeleton organization followed closely with a strength of 1.34 (Figure 3f). Molecular functions associated with the spinal cord hits, such as MHC class I protein binding and structural constituent of cytoskeleton, exhibited strong enrichments with strengths of 2.05 and 1.97, respectively (Figure 3g). Additional functions included GTPase activity and binding, microtubule binding, and various ribonucleotide binding functions. The inclusion of metabolic proteins, particularly GAPDH, and the emphasis on nitrosylation pathways, contribute novel insights to the understanding of cytoskeletal dynamics and synaptic plasticity in the context of SOD1 trimer binding in the spinal cord. The shared pathways from the brain and spinal cord interactome converge to highlight a potentially shared mechanism of SOD1 trimers in the CNS.

### SOD1 trimer skeletal muscle interactome reveals protein biosynthetic process, glycolytic processes, and overall muscle cell homeostasis

Muscle wasting is a clinical hallmark of ALS. The debate surrounding whether this wasting results solely from neuronal degradation or if there exists a pathology originating within the muscle that influences the associated neurons has fueled our motivation to explore the skeletal muscle SOD1 trimer interactome. Our objective was to uncover proteins and pathways unique to the muscle context, shedding light on the potential intrinsic factors contributing to muscle wasting in ALS. We mapped the PPI specific to skeletal muscle and revealed a distinctive landscape encompassing metabolic proteins and those crucial for muscle cell homeostasis (Figure 4a). Central nodes in this interactome included Aconitase 2 (ACO2), aldolase fructose-bisphosphate aldolase A (ALDOA), GAPDH, and triosephosphate isomerase 1 (TPI1). Fructose-bisphosphate aldolase A, exhibiting a fold change of 2.1 with a p-value of 0.0001, emerged as the most significant hit within the skeletal muscle interactome (Figure 4b). We identified a total of 63 identified proteins from the skeletal muscle lysate pulldowns displaying a normal distribution of fold change (Figure 4c). Notably, metabolic proteins—ALDOA, GAPDH, TPI1, and ACO2— represented the main clusters, along with several other proteins lacking known interactions from curated databases in STRING.db or experimental determinations related to the mentioned metabolic proteins (Figure 4e). Biological processes associated with the skeletal muscle hits encompassed the methylglyoxal biosynthetic process (strength 2.87), muscle cell homeostasis (strength 2.35), glycolytic process (strength 2.08), and aerobic respiration (strength 1.84) (Figure 4f). The sole cellular component resulting from the skeletal muscle hits was the myelin sheath, with a strength of 1.57 (Figure 4g).

**Figure 4.**
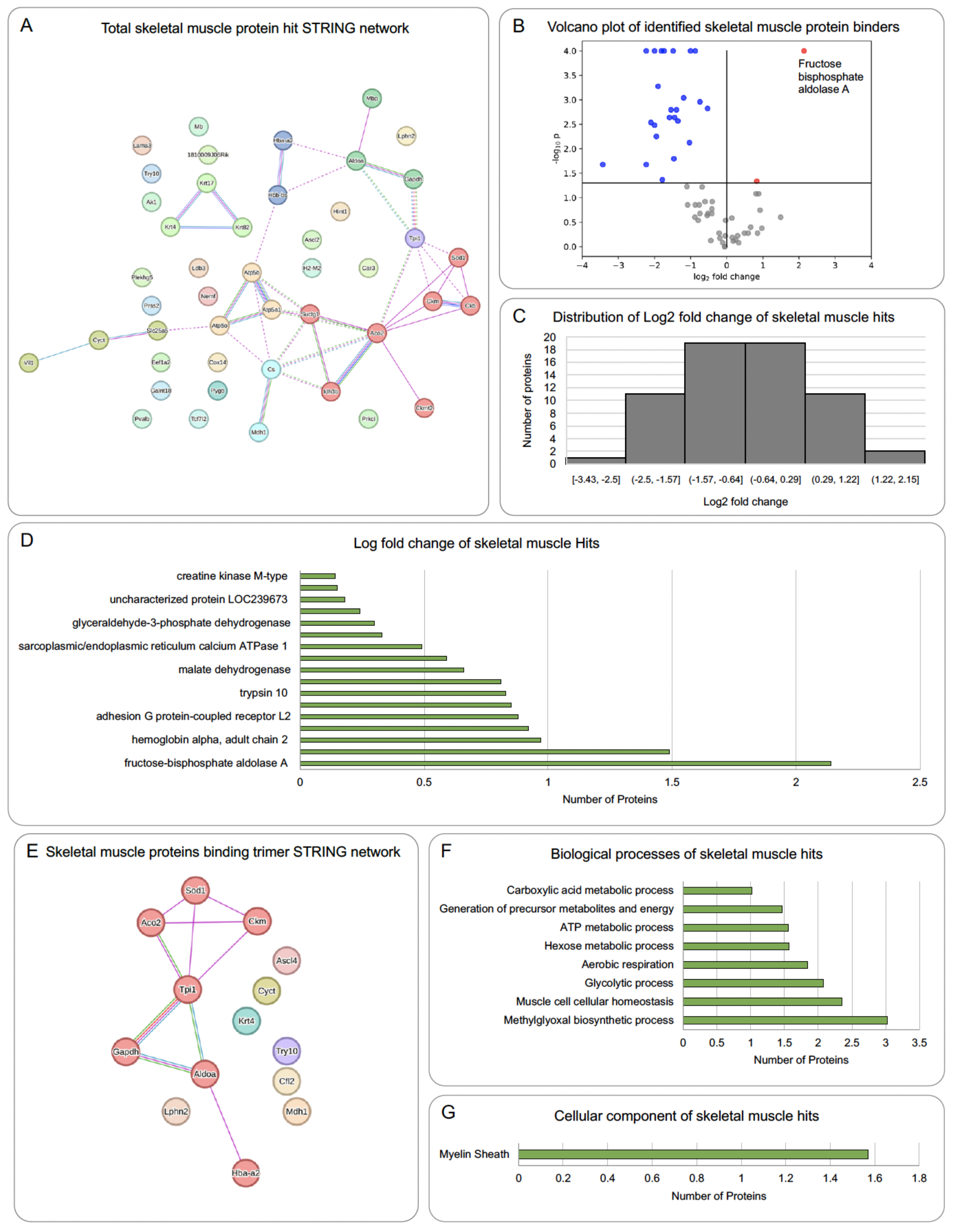
SOD1 trimer skeletal muscle interactome. A) Protein-protein interaction network of SOD1 trimer hits with WT mouse skeletal muscle tissue. B). Volcano plot of SOD1 trimer skeletal muscle hits. Blue are positive fold change nodes and red are negative fold change nodes. Fructose bisphosphate aldolase A was the only hit of statistical significance (p=<0.001. Mann-Whitney test). C). Distribution of fold change of skeletal muscle hits showing normal distribution. D) Fold change of all the skeletal muscle hits ordered from least change to the most change at the bottom. E) Protein-protein network of positive fold change hits displaying one main cluster in red with metabolic proteins centralized. F) Biological processes of the skeletal muscle hits ordered by strength. G) Cellular component of skeletal muscle hits to the myelin sheath.

A striking observation was the prominence of metabolic proteins in the skeletal muscle hits compared to other CNS tissues: the brain and spinal cord. This divergence implies that intrinsic errors in muscle cell homeostasis may contribute significantly to the overall disease process, challenging the assumption that muscle consequences are solely a result of neuron degeneration and CNS dysfunction (Table 1). By identifying unique proteins and pathways specific to the muscle context, we contribute to the evolving understanding of ALS pathogenesis, emphasizing the potential role of intrinsic muscle cell homeostatic errors in the disease process.

### NSC-34 cells expressing SOD1 trimers have differential expression patterns compared to NSC-34 cells expressing SOD1 dimer only and support pathways identified in the interactome

Our interest extended to the investigation of the SOD1 trimer transcriptome as an orthogonal experiment complementing the interactome studies. To discern the distinct effects of various SOD1 trimer mutants, including a destabilized mutant (D101I), a positive control (A4V), and two super stable trimer groups (F20L-H46Q (FH) and H46G-G108H (HG))^22,25,39^, we performed assessments within the Reactome^40^ framework (Figure 5a). The Reactome serves as a comprehensive database encompassing reactions, pathways, and biological processes. We performed differential gene expression analysis from the RNA-seq data and found several genes that are unique to each condition as well as shared between the different groups (Figure 5b). In total, we detected 11360 genes with 16 genes shared exclusively among the trimer groups. We generated volcano plots to illustrate the differentially regulated genes for each trimer (FH and HG) compared to WT, as well as the destabilizing trimer (D101I) compared to WT after 24 hours of trimer expression. Notably, 12 genes were up- regulated and 40 down-regulated in the FH sample, while 22 genes were up-regulated and 45 down- regulated in the HG sample compared to WT (Figure 5b).

**Figure 5.**
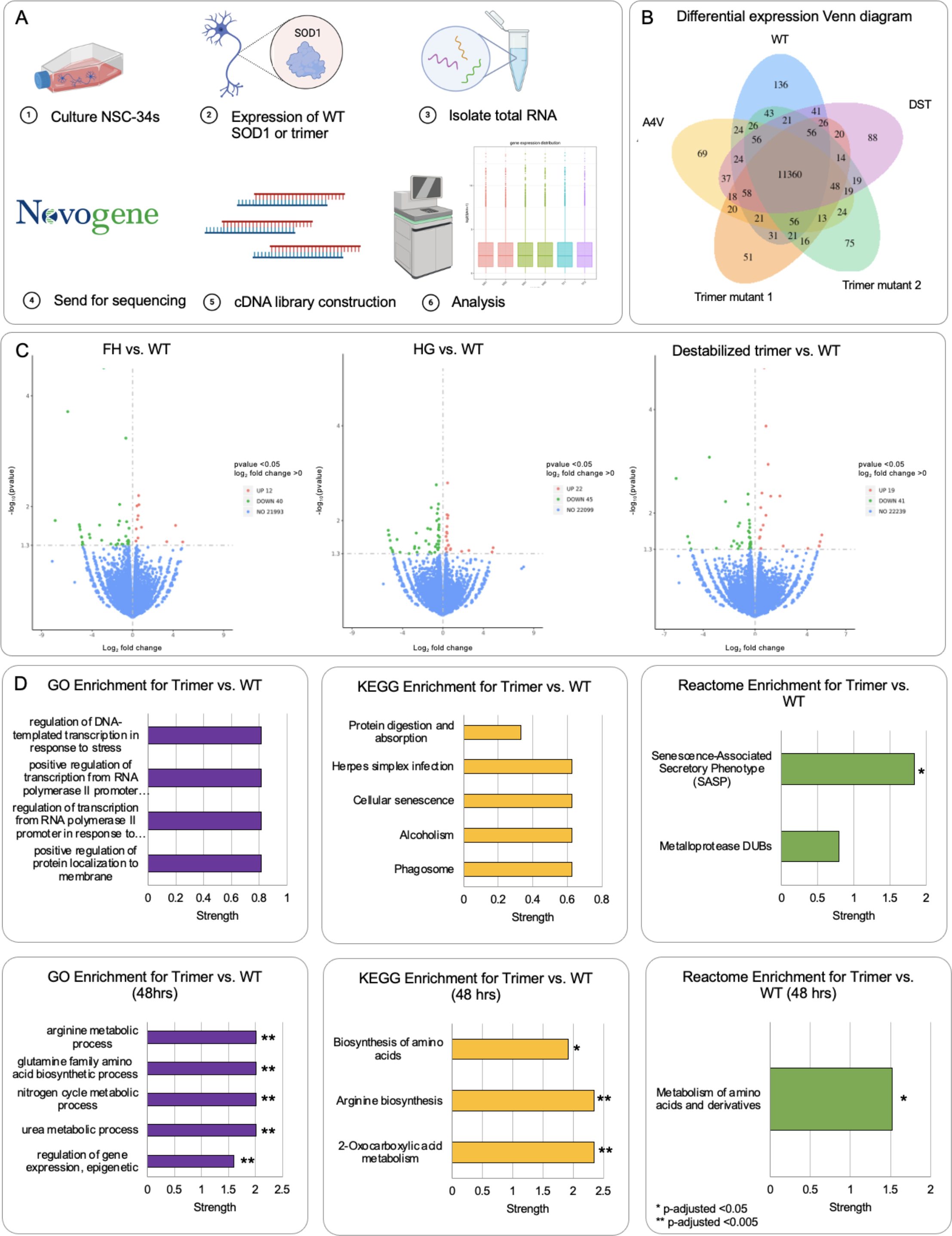
SOD1 trimer transcriptome. A) Methodology schematic for collecting RNA from NSC-34 cells transfected with WT or trimer SOD1 and sequencing for the generation of the SOD1 trimer transcriptome (created with BioRender.com). B) 5-way Venn diagram of differential gene expression for WT, A4V positive control, FH trimer, HG trimer, and destabilizing trimer negative control (DST) for 24-hour expression. C) Volcano plots for differential expression of WT SOD1 and trimer stabilizing mutants and trimer destabilizing. D) GO functions for trimer vs. WT, KEGG pathways for trimer vs. WT, and Reactome pathways for trimers vs. WT at 24-hour and 48-hour expression timepoints. The enrichment analysis was performed using ClusterProfiler36 (v.3.8.1) with a *p-adjusted <0.05 **p- adjusted <0.005.

We performed functional analysis of the RNA-seq data and reported Gene Ontology (GO)^41,42^, KEGG^43^, and Reactome enrichment hits for trimer compared to WT after 24- and 48-hour expression of trimers in NSC-34 cells (Figure 5d). After 24 hours, GO enrichment suggested trends in positive regulation of protein localization to the membrane, regulation of transcription from RNA polymerase II promoter in response to stress, and regulation of DNA-templated transcription in response to stress. Significant GO enrichment pathways after 48 hours included epigenetic regulation of gene expression, urea metabolic process, nitrogen cycle metabolic process, glutamine family amino acid biosynthetic process, and arginine metabolic process (p-adj < 0.005). Top KEGG pathways attributed to FH trimer mutant after 24- and 48-hours indicated trends in phagosome, cellular senescence, and protein digestion and absorption after 24 hours. Statistically significant KEGG pathways after 48 hours included biosynthesis of amino acids, 2-oxocarboxylic acid metabolism, and arginine biosynthesis (p < 0.005). In Reactome enrichment, the Senescence-Associated Secretory Pathway (SASP) was enriched (p < 0.05) after 24- hour trimer expression with a fold change of 1.83. At 48 hours, the metabolism of amino acids and derivatives Reactome pathway was significant with a fold change of 1.52 (p < 0.05). Our transcriptomic studies of NSC-34 cells expressing SOD1 trimers or dimers at 24- and 48-hour time points revealed distinct differences in gene expression, indicating the overall activation of cellular senescence phenotypes and stress responses. Our comprehensive analysis contributes to our understanding of the intricate molecular mechanisms associated with SOD1 trimer mutants, shedding light on potential implications for cellular senescence and stress response pathways.

### Septin-7 binds SOD1 trimers and not native state SOD1 dimers

Septin-7 emerged has a potential binder of SOD1 trimers from the brain and spinal cord hits. Septin-7 is a highly conserved GTP-binding protein and known to play a role in neuronal processes such as regulation of axons and dendrites, synaptic plasticity, and vesicular trafficking. We validated the interaction between SOD1 trimer and septin-7. We combined computational simulations of the complex using discrete molecular dynamics (DMD) with subsequent validation using biochemical experiments that included microscale thermophoresis (MST)^44,45^ and confocal imaging. To perform DMD simulations, we retrieved the septin-7 protein structure (PDB number: 6N0B) from the PDB database and employed a SOD1 trimer model (structure 9) from Proctor et al.^22^ Utilizing ClusPro^46^, we docked septin-7 and the SOD1 trimer model, resulting in ten predicted binding sites (Figure S1). Subsequently, DMD simulations (Figure S3) allowed us to predict the likely residues participating in the binding interaction between the alpha helix on septin-7 (residues 29-47) and parts of the *C* chain on the SOD1 trimer (residues 101-131, 150-154) (Figure 6a).

**Figure 6.**
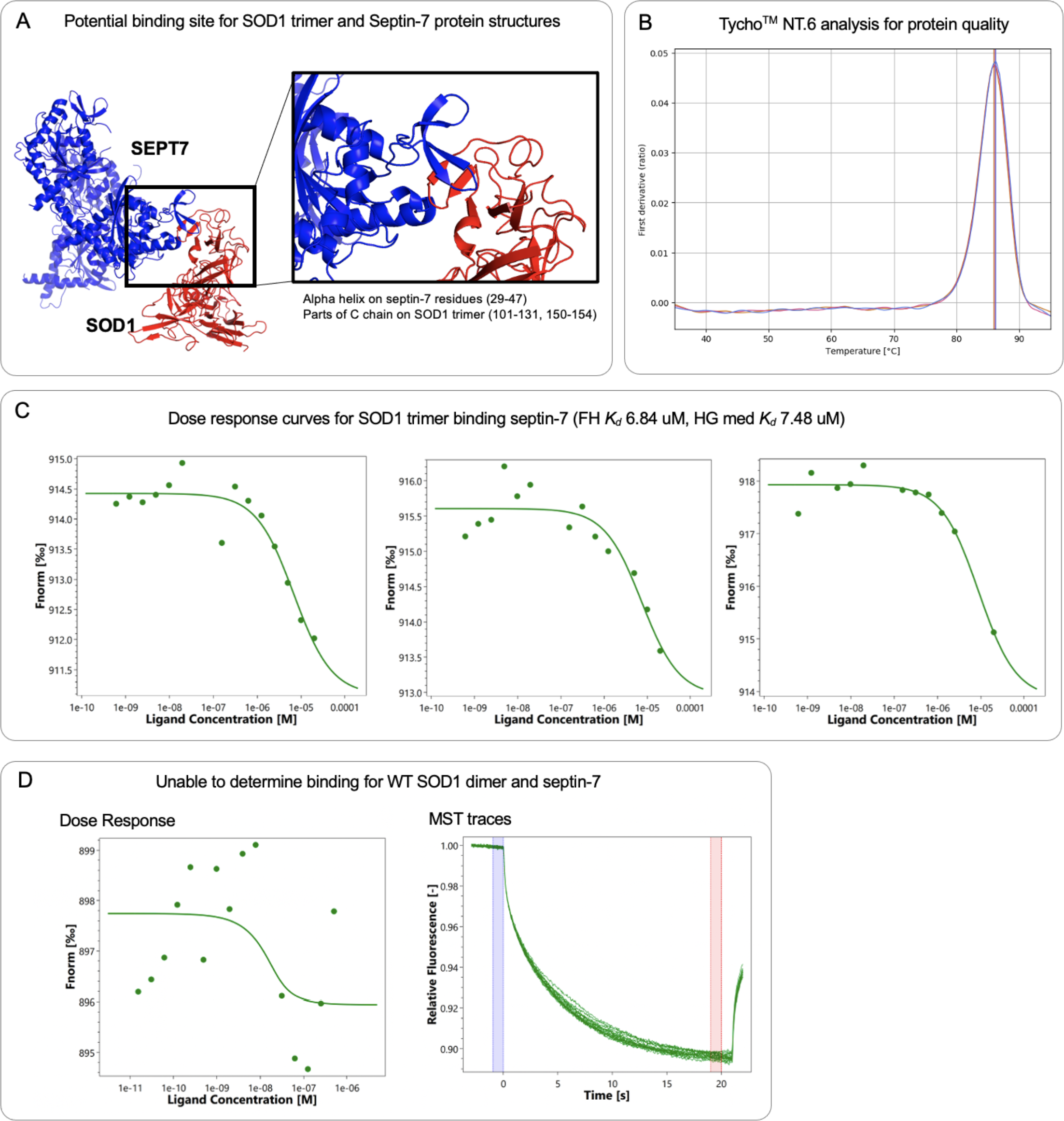
SOD1 and Septin-7 trimer binding validation. A) ClusPro 2.0 docking of SOD1 and Septin- 7 protein structures show a predicted binding site. The predicted binding site includes the alpha helix on septin-7 residues (249-47) and parts of the C chain on SOD1 trimer residues (101-131, 150-154). C) Tycho NT.6 graph showing protein quality and stability of Septin-7 used for subsequent microscale thermophoresis experiments (MST). C) Dose response curves for MST experiments testing binding between SOD1 trimer and septin-7. Trimer mutant bound septin-7 with a kD of 6.84 uM. HG trimer mutant had a median kD of 7.48 uM binding septin-7. D) Dose response curves of MST experiments of WT SOD1 dimer and septin-7. There was no binding detectable with normal MST traces.

To experimentally validate the binding between SOD1 trimers and septin-7, we purified SOD1 trimers, ensuring their protein quality and stability (Figure S2). We obtained purified septin-7 commercially and subjected the proteins (MST) (Figure 6b), generating dose-response curves for the protein-protein interaction (Figure 6c). From MST experiments, we calculated the dissociation constant (*Kd* for the SOD1 trimer binding to septin-7, resulting in 6.84 μM for the FH trimer mutant and a median *Kd* of 7.48 μM for the 2nd HG trimer mutant. To contextualize the significance of these binding measurements, we determined the concentrations of SOD1 and septin-7 in the brain using PAXdp ([SEPT7 10uM] and [SOD1 250 μM]). Following estimations from Khare et al.^47^. and Hnath et al^25^ accounting for potential aggregation, we approximated about 8.8 μM of SOD1 trimer bound out of 10 μM septin-7, underscoring the physiological significance of our calculated *Kd* from MST (Figure S4). In contrast, MST experiments with purified WT SOD1 dimer and septin-7 indicated no binding (Figure 6c). This comprehensive approach not only computationally predicted the binding interface between SOD1 trimers and septin-7 but also provided robust biochemical validation through MST. The observed significant binding, especially in comparison to SOD1 dimers, underscores the relevance of the SOD1 trimer-septin-7 interaction. This work advances our understanding of the molecular associations within ALS pathology, particularly highlighting the role of septin-7 in binding with SOD1 trimers, thus contributing to the broader comprehension of disease mechanisms.

### Septin-7 colocalizes with misfolded SOD1 but not native SOD1 dimers

We performed confocal imaging, followed by colocalization analysis, using the C4F6 antibody^48,49^ to stain SOD1 trimers and septin-7 (Figure 7a). The C4F6 antibody specifically binds to misfolded SOD1, including SOD1 trimers^22,25^. Representative images for all control and experimental groups are provided in Figure S5. Notably, we observed C4F6 antibody staining for the presence of misfolded SOD1 across all groups. We observed septin-7 staining diffusely in stably expressing NSC-34 cells, with a notable presence along the outer edges of the cells. We identified positive co-localization of C4F6, and septin-7 compared to WT SOD1 staining (Figure 7b), while no significant co-localization of trimers with nuclei was observed (Figure 7c). Beyond qualitative interpretation, our quantitative measurements of colocalization revealed significant differences between native and misfolded SOD1 for septin-7, but not for Hoechst (Table 2). Anticipating a low misfolded SOD1 signal in the wild-type group, we performed separate analyses for WT SOD1 transfected NSC-34 cells compared to the trimer stabilizing mutants (FH and HG). Three out of five methods detected a significant difference in colocalization with septin-7 between the two types of SOD1 in both stratifications. In contrast, colocalization of SOD1 with Hoechst was only significant for one out of the ten comparisons performed.

**Figure 7.**
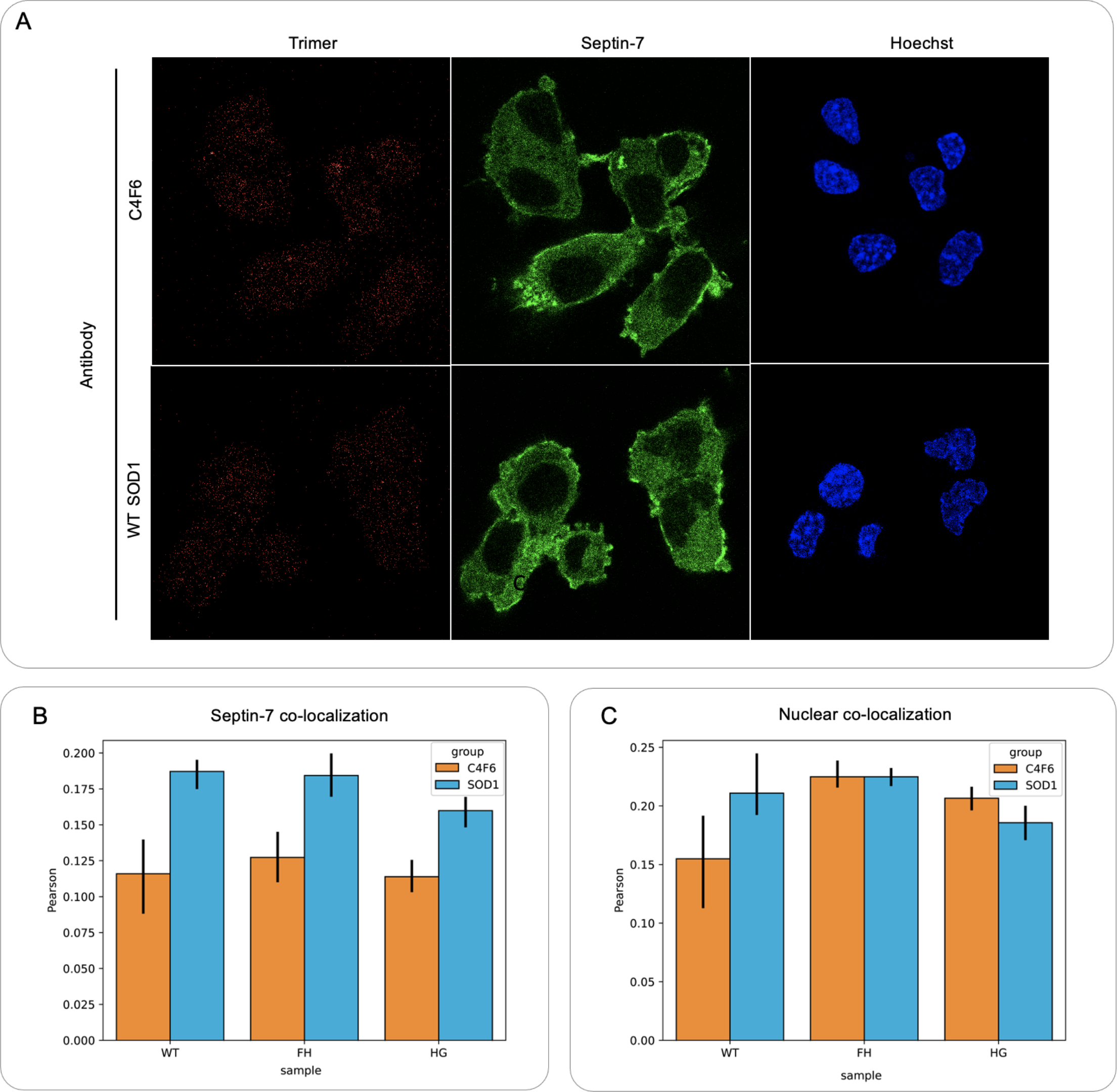
Co-localization of SOD1 trimer and Septin-7 in NSC-34. A) Confocal images of differentiated stably expressing SOD1 in NSC-34 cells stained for septin-7 (green) or SOD1 (red) or nuclei (Hoechst). The top row is C4F6 antibody staining which detects misfolded SOD1 compared to the bottom row which is WT SOD1 dimer antibody staining. B) Septin-7 co-localization with misfolded SOD1 (C4F6) and WT SOD1 compared between different cell lines (WT, FH, and HG trimer mutants). D) Nuclear co-localization with misfolded SOD1 (C4F6) and WT SOD1 compared between different cell lines (WT, FH, and HG trimer mutants).

**Table 2:** Results of Welch’s t-test for unequal variances, testing for differences in colocalization of native vs misfolded 50D1 with septin-7 and Hoechst.

| **Colocalization target** | **Sample** | **t (p-value) for native vs misfolded SOD1** | **degrees of freedom** |
| --- | --- | --- | --- |
| Septin-7 | WT | 5.76 (0.0003)* | 8.18 |
|  | Trimer | 5.02 (< 0.0001)* | 31.9 |
| Hoechst | WT | 0.511 (0.64) | 2.82 |
|  | Trimer | 0.695 (0.49) | 33.5 |

We employed the Costes’ randomization method to validate colocalization, revealing significant colocalization at the p = 0.05 level for 34 out of 38 (89%) of the septin-7 colocalizations and 35 out of 37 (94%) of the Hoechst colocalizations (Table S1). Our colocalization analysis provides compelling evidence for the association between SOD1 trimers and septin-7, reinforcing the significance of their interaction. These findings enhance our understanding of the spatial relationships within NSC-34 cells, shedding light on the potential implications of misfolded SOD1 and its colocalization with septin-7.

## Materials and Methods

### Mouse tissue lysates

We obtained 17-month-old wild-type black CD-1 male mice (RRID:IMSR_CRL:022) from the Penn State Animal Research Facility (ARF). The ARF performed cervical dislocation to sacrifice the mice and directly after, we dissected the brain, spinal cord, and tibialis anterior muscle from each mouse, and froze the tissues in liquid nitrogen. We combined 0.4 mL of RIPA lysis buffer with phosphatase and protease inhibitors per 0.1 g of tissue. We placed the resuspended tissue in a glass homogenizer for manual homogenization. Then, after manual homogenization, we incubated the homogenate on ice for 10 minutes and centrifuged the lysate at 19,000 g for 10 minutes at 4 °C. We collected the supernatant and saved the protein tissue lysate for bicinchoninic acid assay (BCA) protein concentration analysis^50^ as well as SDS page gel analysis with Coomassie blue staining for visualization of the collected tissue protein lysates.

### Preparation of the carboxyl beads

We suspended carboxyl magnetic beads (AbraMag^TM^) in coupling buffer (0.1 M 2-(N-morpholino) ethanesulfonic acid (MES) pH 6.0) and subsequently washed the beads in the buffer. We added 250 uL of 1-ethyl-3-(3-dimethylaminopropyl) carbodiimide hydrochloride/N-Hydroxysuccinimide (4 mg EDC/0.6 mgNHS) solution and rotated the beads at room temperature for 15 minutes. Then, we performed three additional wash steps in the coupling buffer before adding 1.5 mg of purified SOD1 to 250 uL of beads (WT and trimers (F20L-H46Q (FH) and H46G-G108H (HG))^22,25,39^ purified using previously described yeast purification methods and size exclusion chromatography^22,25^). The purified proteins were tested for correct folding through circular dichroism^51,52^ and with size exclusion chromatography and silver-stained SDS page gel. We rotated the mixture overnight at room temperature to facilitate SOD1 protein binding to the carboxyl magnetic beads. The next morning, we removed the supernatant and calculated the percent of protein bound by measuring the optical density at 280 nm and comparing it to the initial amount of protein added. We added carboxyl quenching buffer (50 nM Ethanolamine pH 8 0.1 % BSA) and rotated the beads at room temperature for 30 minutes. After quenching, we washed the SOD1 coupled carboxyl magnetic beads with PBS and stored in PBST at 4 °C. We generated WT dimeric SOD1, and two SOD1 trimers (F20L-H46Q (FH) and H46Q-G108H (HG) mutations) coupled sets of magnetic beads for the pulldown assays.

### Pulldown assays

We washed the carboxyl magnetic beads, coupled with SOD1 (WT, FH, HG), in PBS then resuspended the beads in each tissue lysate (brain, muscle, or spinal cord) after optimizing the lysate concentration. We facilitated the binding of SOD1 and tissue proteins overnight at 37 °C. Then, we used a magnet to sequester the beads and removed unbound extraneous protein. We washed the beads three times with PBS and then eluted the bound protein with elution buffer (0.1 M citric acid pH 2.5). After elution, we stabilized the eluted protein by adding 1 M Tris pH 9 until the solution pH reached a physiological pH of 7.4. We washed the beads with PBS and repeated the pulldowns with WT, FH, and HG coupled beads with brain, muscle, and spinal cord tissue lysates in triplicate.

### iTRAQ labeling and mass spectrometry analysis

After we performed the pulldown assays, we collected the protein that bound to WT, FH, and HG SOD1 in the brain, muscle, and spinal cord with three replicates and lyophilized the samples. To each sample, we added triethylammonium bicarbonate buffer (TEAB) and 2 % SDS to denature the protein sample.

We added the reducing reagent (100 mM TCEP) to each sample then we incubated the samples at 60 °C for 1 hour. We centrifuged the samples and added 84 mM iodoacetamide and incubated the tubes in the dark at room temperature for 30 minutes. We prepared mass spectrometry grade trypsin with 50 mM acetic acid resuspension buffer and added 1 mg/mL solution of trypsin to each sample. We incubated the samples with trypsin at 48 °C overnight and lyophilized the sample if the volume exceeded more than 33 uL. We prepared the iTRAQ^53,54^ label according to manufacturer’s instructions and added the iTRAQ labels to each sample for incubation at room temperature for 2 hours. We then quenched the iTRAQ label reaction with molecular biology grade water at room temperature for 30 minutes and dried the sample completely. We quenched and dried the samples each three more times before bringing the samples to the Penn State College of Medicine Proteomics and Mass Spectrometry core facility for mass spectrometry (MS) analysis. The mass spectrometry proteomics data have been deposited to the ProteomeXchange Consortium^55^ via the PRIDE^56^ partner repository with the dataset identifier PXD050873.

The Penn State College of Medicine Proteomics and Mass Spectrometry core facility used the Scaffold Q+ (version Scaffold_5.0.1, Proteome Software Inc., Portland, OR) to quantitate Label Based Quantitation (iTRAQ) peptide and protein identifications. They accepted peptide identifications if they could be established at greater than 95.0% probability by the Scaffold Local FDR algorithm. Protein identifications were accepted if they could be established at greater than 99.0 % probability and contained at least 2 identified peptides. They assigned protein probabilities by the Protein Prophet algorithm^57^. If there were proteins that contained similar peptides and they could not differentiate based on MS/MS analysis alone to satisfy the principles of parsimony. They grouped proteins sharing significant peptide evidence into clusters.

They corrected channels by the matrix:

[0.000,0.000,0.929,0.0689,0.00240]; [0.000,0.00940,0.930,0.0590,0.00160];

[0.000,0.0188,0.931,0.0490,0.001000];[0.000,0.0282,0.932,0.0390,0.000700];

[0.000600,0.0377,0.933,0.0288,0.000];[0.000900,0.0471,0.933,0.0191,0.000];

[0.00140,0.0566,0.933,0.00870,0.000]; [0.00270,0.0744,0.921,0.00180,0.000]

in all samples according to the algorithm described in i-Tracker (Shadforth, I et al BMC Genomics 2005;6 145-151). They performed normalization iteratively (across samples and spectra) on intensities, as described in Statistical Analysis of Relative Labeled Mass Spectrometry Data from Complex Samples Using ANOVA^58^. They used medians for averaging then they log-transformed the spectra data, pruned of those matching to multiple proteins, and weighted by an adaptive intensity weighting algorithm. Of 1983 spectra in the experiment at the given thresholds, 1480 (75 %) were included in quantitation. Finally, they determined differentially expressed proteins by applying Mann-Whitney Test with unadjusted significance level p < 0.05 corrected by Benjamini-Hochberg. We used STRING.db^30,31^ to visualize the protein-protein interaction networks.

### RNA-seq of SOD1 trimers in NSC-34 cells

We cultured differentiated NSC-34 cells (RRID:CVCL_D356) (differentiation media DMEM:F12 10 % FBS, 1 % L-Glutamate, 1 % Penn/Strep, 0.5 % non-essential amino acids, 10 mM Retinoic acid) over 3 days and performed a transfection (jetOPTIMUS) with SOD1 WT and trimer constructs for expression. We collected the cell lysates with RIPA buffer at 24- and 48-hour timepoints post-transfection. RNA was extracted using the RNeasy mini kit (Qiagen) and total RNA concentration was determined using NanoDrop OD readings at 260/280 wavelengths, the samples were then sent for RNA-seq analysis (Novogene, Beijing, China). Novogene quality checked the raw data with error rate distribution, GC-content, and data filtering. Then, the data was mapped to the reference genome and gene expression was quantified. Differential analysis of the RNA-seq data was performed using DESeq2^59^ (v1.26.0) with a p-adjusted value of ≤ 0.05. Functional analysis included enrichment analysis, oncogene analysis and protein-protein interaction analysis. The enrichment analysis was performed using ClusterProfiler^60^ (v.3.8.1) with a p-adjusted value < 0.05.

### Molecular modeling of SOD1 trimers in complex with interactor proteins

Proctor et al. has developed an SOD1 trimer model^22^ used in this study, and we used available structures of interactor proteins (septin-7 PDB ID: 6N0B). We removed clashes using Gaia^61^ and Chiron^62^ and used ClusPro^46^ to predict protein-protein binding. We performed docking of SOD1 trimer against the interactor proteins to reveal putative binding sites. For illustrative purposes, we used discrete molecular dynamics^63–65^ to simulate one of the docked poses for the SOD1 trimer and the Septin-7 dimer. We overlaid this simulation with the Septin-2G/Septin-6/Septin-7 hetero-hexamer (PDB ID 7M6J).

### Microscale thermophoresis to characterize binding interactions

We performed microscale thermophoresis^44,66^ (MST) to quantify the binding interactions. We purchased commercially purified Septin-7 (LSBio cat# LS-G23386-100) and used purified SOD1 WT and trimer as described by Hnath and Dokholyan^25^. Then, we performed MST studies (Monolith) by first, labeling our target (septin-7) with the Monolith His-Tag labeling kit (RED-tris-NTA 2^nd^ generation). We performed pre-tests to evaluate the fluorescence intensity and homogeneity. We assessed the formation of aggregates and adsorption. Next, we performed a binding check and determined the optimal assay conditions such as buffer (PBS) and binding affinity. The MST graphs calculated the Kd and we followed our initial experiments with assay optimization.

### Generation of NSC-34 cells stably expressing SOD1

We generated NSC-34 stable cells expressing SOD1 (WT, A4V or trimers) using the Piggybac transposon transduction protocol^67^. SOD1 or a GFP control was cloned into the dox-XLone plasmid. The NSC-34 cells were transfected with dox-XLone plasmid with SOD1 or GFP with the pPB-UbC (PiggyBac) plasmid 1:1 using JetOPTIMUS reagents. We determined the appropriate concentration of Blasticin for the selection of successfully transfected cells and performed selection. After selection, we amplified and froze cell line stocks at -80 °C. For experiments, we continued blasticidin (10 ug) selection and differentiated the stable cell lines by adding 10 μM retinoic acid, we induced expression of GFP or SOD1 using doxycycline at 2 ug/mL. We confirmed protein expression with fluorescence of GFP in the control group and by western blot and then native gels for SOD1 trimers.

### Confocal imaging of NSC-34 cells expressing SOD1 trimers

We cultured differentiated NSC-34 cells, stably expressing WT, controls (A4V and destabilizing trimer mutant D101I), and super stable trimers, at 5,000 cells/well on 35 mm glass bottom dishes coated with ESGRO complete gelatin solution (Millipore) for 3 days. Then, we fixed the cells with 4 % paraformaldehyde and permeabilized the cells for immunostaining with 0.05 % triton-X100. We added the primary antibodies rabbit anti-septin-7 (RRID:AB_2647129, Thermo Fisher Scientific Cat# PA5- 56181) and mouse anti-C4F6 SOD1 antibody^68^ (MediMabs Cat# MM-0070-2-P, RRID:AB_10015296). We then added respective secondary antibodies Alexa Fluor 488 goat anti-rabbit (Thermo Fisher Scientific Cat# R37116, RRID:AB_2556544) and Alexa Fluor 635 goat anti-mouse (Thermo Fisher Scientific Cat# A-31574, RRID:AB_2536184) for visualization of primary antibodies. We then added Hoechst 33342 (Invitrogen Cat# H3570) for 10 minutes for nuclear staining. For the confocal imaging, we used the Leica SP8 STED 3x microscope at the Penn State College of Medicine Advanced Light Microscopy core facility. We processed and analyzed the images using the LAS X Life Science Microscope software.

### Quantification of colocalization

We used Fiji and the JACoP plugin to quantify 2D colocalization of (1) septin-7 and (2) Hoechst with both native and misfolded SOD1. We report on the following metrics of colocalization: Pearson correlation coefficient (PCC), Costes’ automatically thresholded PCC, Manders’ overlap coefficient (with manually chosen thresholds of 100 for Hoechst, 1 for native and misfolded SOD1, and 50 for septin-7), Van Steesel’s cross-correlation, Li’s intensity correlation quotient, and Costes’ randomization approach. To quantify differences in colocalization among the various groups, we performed Welch’s t-test for unequal variances using SciPy. For the PCC and Costes’ PCC, we first performed a Fisher r-to-z transform.

### Western blot analysis

NSC-34 stable cell lines expressing SOD1 (WT or trimer) lysate from 3 wells of a six-well plate are used for each sample. We collected the lysates from the wells using RIPA buffer (with phosphatase and protease inhibitors from Pierce VWR) and scraping the wells. We centrifuged the samples at 13,000 x g for 10 minutes to pellet the lysed cells. We denatured the lysates and resolve using 10 % (v/v) SDS-PAGE before blotting on PVDF membranes using TransBlot Turbo (BioRad) for transfer. We incubated the blots with antibodies against SOD1 (Sigma Cat# SAB5200083, RRID:AB_2890259) or C4F6 (MediMabs Cat# MM-0070-2-P, RRID:AB_10015296) followed by secondary antibody conjugated to HRP (Thermo Fisher Scientific Cat# A16096, RRID:AB_2534770). We visualized the blots using a ChemiDoc MP system (BioRad).

## Discussion

The complexity of ALS etiology poses a significant challenge for researchers and clinicians seeking to identify crucial, disease-modifying nodes within the intricate network of cell pathways^17,69^. Compounding this challenge is the heterogeneous nature of ALS, with both sporadic and familial forms exhibiting a spectrum of genetic mutations. Our investigation into the direct protein interactions of SOD1 trimers and subsequent pathway analyses link seemingly disparate genes and mechanisms associated with ALS and the continuum of broader neurodegenerative disorders. The presence of unique hits in each tissue type suggests that intrinsic SOD1 trimers may induce disruption specific to each system, contributing to the overall disease complexity rather than originating from a singular source. The degeneration of neurons affects several different tissue types such as the brain, spinal cord, and skeletal muscle. Muscle wasting and loss is a clinical hallmark for ALS patients. The debate^70–72^ regarding whether the disease originates in the muscle or CNS adds an additional layer of complexity. Our findings underscore the importance of proteins influencing cytoskeletal structure and synaptic plasticity in CNS tissues, contrasting with strong metabolic themes observed in SOD1 trimer binding in muscle tissue. Maintaining muscle metabolic homeostasis and synaptic plasticity in neurons are crucial aspects. Importantly, our work highlights tissue-selective and within-cell compartment-selective binding of SOD1 trimers compared to their WT native dimer counterparts. This nuanced understanding contributes to unraveling the intricate landscape of ALS pathology, providing valuable insights for targeted therapeutic interventions.

Interactions involving the SOD1 trimer protein in the brain exhibit a high degree of localization, primarily influencing the organization of neuronal cytoskeletons and the regulation of dendritic spines, thereby impacting synaptic plasticity. Notably, these interactions are heavily influenced by phosphorylation, with the mislocalization of the SOD1 trimer binders which are known to have detrimental effects on neuronal health and communication. Within the CNS hits, microtubule-associated protein tau (MAPT) emerged as a central node in the protein-protein interaction network. MAPT plays a crucial role in promoting microtubule assembly and stability, maintaining neuron polarity, and regulating axonal transport, all while promoting neurite growth^73^. SOD1 trimer binds neurodegenerative disease associated proteins in the brain providing evidence of convergent pathological pathways. Beta-adducin (ADD2), a cytoskeletal junctional complex protein highly expressed in the brain, regulates actin filaments and is a key component of synaptic structures. It influences dendritic spines and processes related to synaptic plasticity, such as learning and memory^74^. ADD2 influences dendritic spines and synaptic plasticity processes such as learning and memory. Another binding partner is brain acid-soluble protein 1 (BASP1), a signaling protein crucial for neurite outgrowth and plasticity. BASP1 exhibits precise localization to synaptic terminals, dendritic spines, and synaptic vesicles, constituting approximately 1% of brain protein during development and around 0.5% in the adult brain. Overexpression of BASP1 in adult neurons stimulates neurite outgrowth^75^. Comparing to other models of ALS, in TDP-43 models, there is a loss of stathmin-2 function^76^. Stathmin-2 regulates the neurofilament-dependent structuring of the axoplasm^77^. This function is critical for maintaining function of neurons. Antisense oligonucleotide-mediated suppression of stathmin-2 in aging mice was able to drive motor neuron death through destruction of axonal caliber and neurofilament spacing^78^. In the spinal cord, the predominant biological process associated with the SOD1 trimer hit was peptidyl-cysteine S-nitrosylation, involving the covalent addition of a nitric oxide group to the sulfur atom of a cysteine residue in a protein. This process implicates mitochondrial proteins and glyceraldehyde-3-phosphate dehydrogenase (GAPDH) and mislocalization of GAPDH to the nucleus, potentially precipitating stress-induced cell death in neurons. Additional noteworthy biological processes include microtubule cytoskeletal organization, mitotic cell cycle, and organelle organization. Our work to understand SOD1 trimer interactions in distinct CNS tissues demonstrates a potential convergence of pathological pathways.

In the skeletal muscle, the protein interactions of the SOD1 trimer predominantly involved metabolic proteins. Notably, fructose-bisphosphate aldolase A emerged as the most significant hit, a protein previously associated with biomarker studies in ALS patients^38^. Aldolase A, a common autoantigen in Alzheimer’s disease and multiple sclerosis^79^ has gained attention as a therapeutic target due to its multifaceted functions in cellular scaffolding, signaling, and motility^80^. Another key player in the skeletal muscle protein-protein interaction network for SOD1 trimers was GAPDH, a glycolytic enzyme crucial for catalyzing the conversion of glyceraldehyde-3-phosphate to 1,3-bisphosphoglycerate. While GAPDH is well-known for its glycolytic role and often utilized as a “housekeeping” protein, it also contributes to RNA export, DNA replication and repair, as well as cytoskeletal organization^81^. Notably, mutant SOD1 expression in ALS is known to decrease S-nitrosylation of mitochondrial proteins and GAPDH^82^. The aberrant gain of function observed in SOD1 in ALS has led to confusion regarding the role of disrupted redox regulation in ALS pathogenesis, and the interaction of SOD1 trimers with proteins involved in S-nitrosylation could offer insights into the causative mechanisms. Hyper-S-nitrosylation of parkin and several other proteins has been observed in the brain tissue of neurodegenerative disease patients^83^. In Alzheimer’s disease, GAPDH interacts with amyloid-beta protein precursor, modulating apoptosis and emerging as a potential therapeutic target. Other significant protein binding partners in the skeletal muscle included mitochondrial aconitase 2 (ACO2), encoding aconitase 2 that catalyzes citrate isomerization in the Krebs cycle. Mutations in ACO2 are associated with cerebellar retinal degeneration and a severe early-onset neurodegenerative condition with seizures^84^. Additionally, MHC class I proteins, known for binding peptides generated from degraded cytosolic proteins, were identified. Neuronal expression of MHC class I is implicated in synaptic plasticity and axonal regeneration in the CNS^85^. The intricate network of these protein interactions in skeletal muscle sheds light on the metabolic and regulatory processes affected by SOD1 trimers. Recent therapeutic efforts from Bravo-Hernandez et al.^86^ showed that spinal delivery of an adeno-associated virus with an shRNA-SOD1 silencing vector after disease onset in an ALS animal models was able to preserve spinal motor neurons as well as preserving muscle innervation. The preservation of the neuromuscular junction from this specific therapeutic may be due to the halting of toxicity from both the nerve and muscle sides.

In parallel, the transcriptomic analysis corroborated the broader patterns identified in the tissue protein binding data. We found enriched cellular pathways related to the regulation of protein localization, metabolic processes, and overarching senescence-associated phenotypes associated with SOD1 trimers when compared to native SOD1 dimers over time. Although variations were observed between the two groups, we identified consistent thematic pathways echoing the protein binding hits, emphasizing an overall impact on cell health maintenance. Our analysis revealed alterations in cellular senescence, representing the cellular aging process, with genes indicating changes attributable to SOD1 trimers in contrast to cells expressing WT dimers. Importantly, the cellular focus of the transcriptomic work allowed us to observe changes at the cellular level that reflect the initial findings from tissue studies.

Following the exploration of our various CNS and muscle binding proteins, we undertook an integrated approach involving computational and biochemical methods to validate the binding of septin-7 with SOD1 trimers specifically in neurons. Septin-7 is a member of the septin protein family, characterized by highly conserved cytoskeletal GTPases. It holds a unique position within the core unit, encompassing septin-6, -7, -2, and -9, and concurrently plays a vital role in dendritic spine morphogenesis and dendrite outgrowth in neurons. The phosphorylation of septin-7, mediated by TAOK2, is instrumental in regulating the formation of mislocalized synapses^87^. Septin-7 has established interactions with HDAC6, contributing to decreased microtubule stability and the regulation of neuron polarity in select neurons. Its significance extends to Alzheimer’s disease, where varying p25 levels, a cleaved component of a tau cyclin-dependent kinase, influence neurodegeneration and memory formation^88^. In neurons, septin-7 assumes a pivotal role in the formation and maintenance of the axon initial segment (AIS), a specialized region at the beginning of the axon where action potentials are generated. Septin-7 contributes to the formation of the cytoskeletal scaffold at the AIS, crucial for regulating neuron excitability and preserving the functional polarity of neurons. Beyond its structural role, septin-7 also governs dendritic spine morphology and synaptic function. While the exact mechanism by which septin-7 exerts its effects in neurons remains unknown, its dynamic involvement in cell structure organization and neuronal function positions it as a compelling target for investigating SOD1 trimer binding. Our validation studies, employing microscale thermophoresis (MST) and colocalization analysis from confocal imaging following pulldown assays, confirmed selective binding of SOD1 trimers compared to native SOD1 dimers. Additionally, we employed molecular docking to predict potential binding sites for future investigations. Disrupting this binding interaction may offer a potential mechanism to mitigate neuronal death, given the known toxicity of SOD1 trimers.

Discontinuity between the complex molecular etiology and phenotypic presentation of in ALS results begs the question whether there is a singular pathway leading to motor neuron death (convergence^89–91^) or the result of distinct and even independent genetic, epigenetic, and environmental pathways (divergence^92–94)^. Our study expands upon prior research on ALS pathophysiology, revealing evidence for a convergence phenomenon centered around SOD1 trimers. We identify SOD1 trimers as pivotal connectors within ALS, elucidating their interactions with other genetically and physiologically linked genes in interactome and reactome maps across different tissue systems. This centralized role challenges divergence models of ALS^95,96^, highlighting overlaps in genes, proteins, and connections to broader neurodegenerative disorders.

## Conclusion

The pathophysiology and mechanisms underlying ALS is complex with numerous disease-modifying mutations that underlie both sporadic and familial cases. Misfolded soluble species of SOD1 are associated with disease severity, and specifically, trimeric forms of SOD1 are toxic compared to larger insoluble aggregates. The role of toxic SOD1 trimers in ALS is unknown. In our study, we constructed SOD1 trimer interactomes from mouse brain, spinal cord, and skeletal muscle tissues. We found tissue-selective SOD1 trimer-protein interactions and subsequent pathway analyses revealed pathways that have previously linked to ALS. In the nerves, we discovered protein interactions that are involved in dendritic spine morphogenesis and synaptic function. In the skeletal muscle, we found proteins that are responsible for maintaining energy and amino acid metabolism. The SOD1 trimer transcriptome from neuron-like cells revealed enrichment of genes associated with the regulation of protein localization, metabolic processes, and overarching senescence-associated phenotypes. Additionally, we found that septin-7 selectively binds SOD1 trimers but not the native dimer through integrative computational and biochemical analyses. Our focus on the early events stemming from SOD1 trimer binding provides a foundation for deciphering critical mechanisms involved in mediating dysfunction and eventual cell death in both neurons and muscle cells. Given the heterogeneity of ALS, we discuss a double-edged sword theory for SOD1 trimer toxicity, where SOD1 trimers induce dysfunction in both the CNS and muscle. Through this work, we identify pivotal protein players and pathways in each system and show potential convergence of genetic and pathophysiological disease mechanisms in ALS through SOD1 trimer interactions. Ultimately, achieving a comprehensive understanding of ALS is essential for advancing ALS therapeutics and the broader understanding of other neurodegenerative diseases.

## Acknowledgements

We thank R.N. and E.P. for proofreading the manuscript. We also thank Drs. Bruce Stanley and Anne Stanley from the Penn State Hershey Mass Spectrometry core for the guidance and analysis of the pulldowns. We thank the Penn State Hershey Animal Research Facility for providing the mice tissue. We also thank Novogene for the RNA-seq analysis.

## Author contributions

N.V.D. supervised the project in its entirety. E.S.C. and B.H. conceptualized the project and the methodology. B. H. purified proteins for pulldowns and advised the experimental design. E.S.C. performed the investigation and formal analysis. C. M. S. performed the computational work and the statistical analysis for the co-localization data. E. S. C. wrote the initial draft, and all authors edited the manuscript.

## Data and materials availability

All data needed to evaluate the conclusions are present in the paper and/or the Supplementary Materials.

