## Supplemental Information for "Unveiling the double-edged sword: SOD1 trimers possess tissue-selective toxicity and bind septin-7 in motor neuron-like cells"

Figure S1. ClusPro 2.0 docking of SOD1 and septin-7 protein structures resulted in ten models of protein-protein binding defined by centers of highly populated clusters of low-energy docked structures. SOD1 trimer predicted structure from Proctor et al. 2016 in blue and septin-7 (PDB number 6N0B) in red.

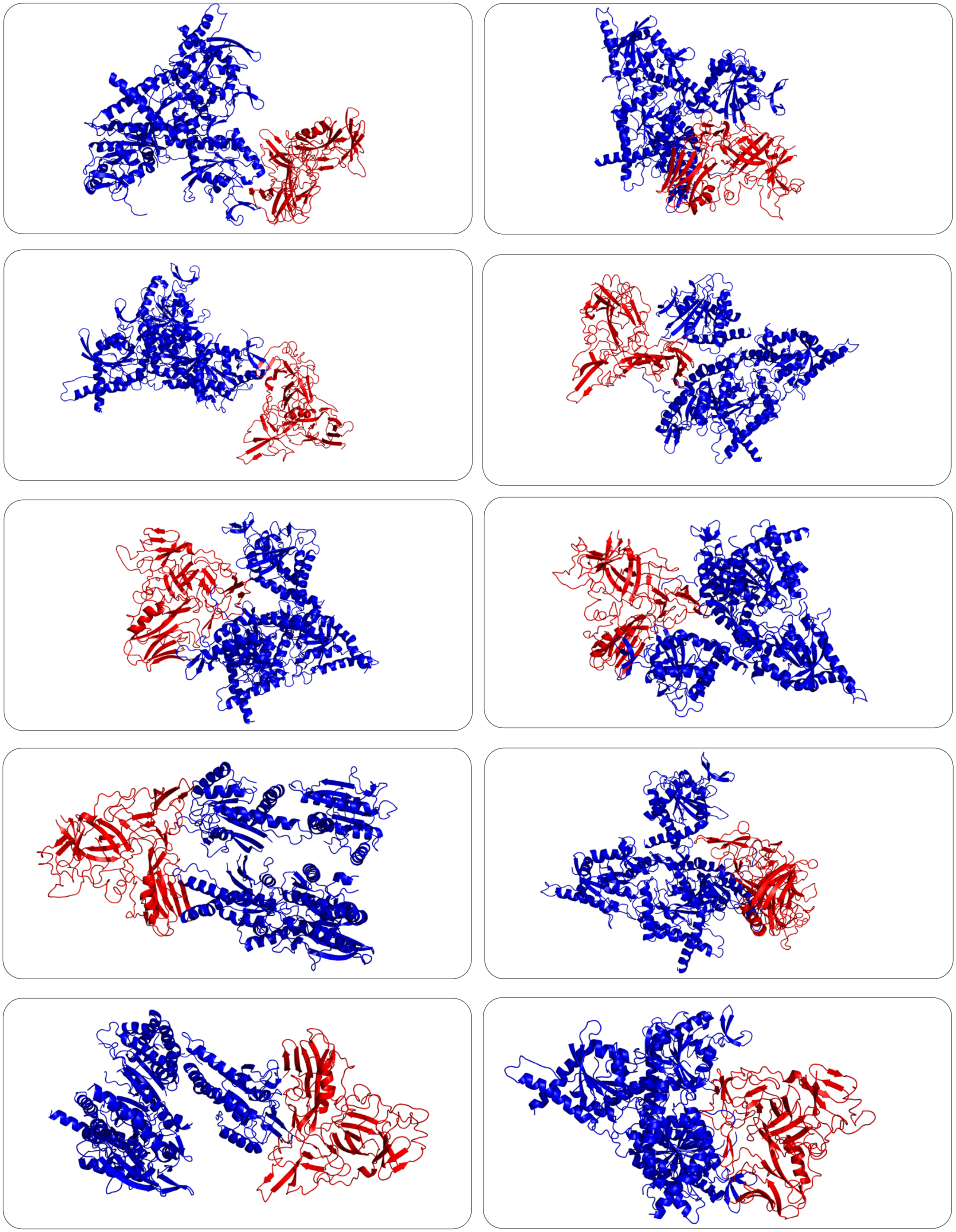

Figure S2. Lab purified SOD1 trimer stability and quality test with Tycho[TM] NT.6 assessment. Trimer mutants (FH and HG) in triplicates for MST experiments. SOD1 trimers were stable and did not denature until 90° Celsius in PBS buffer.

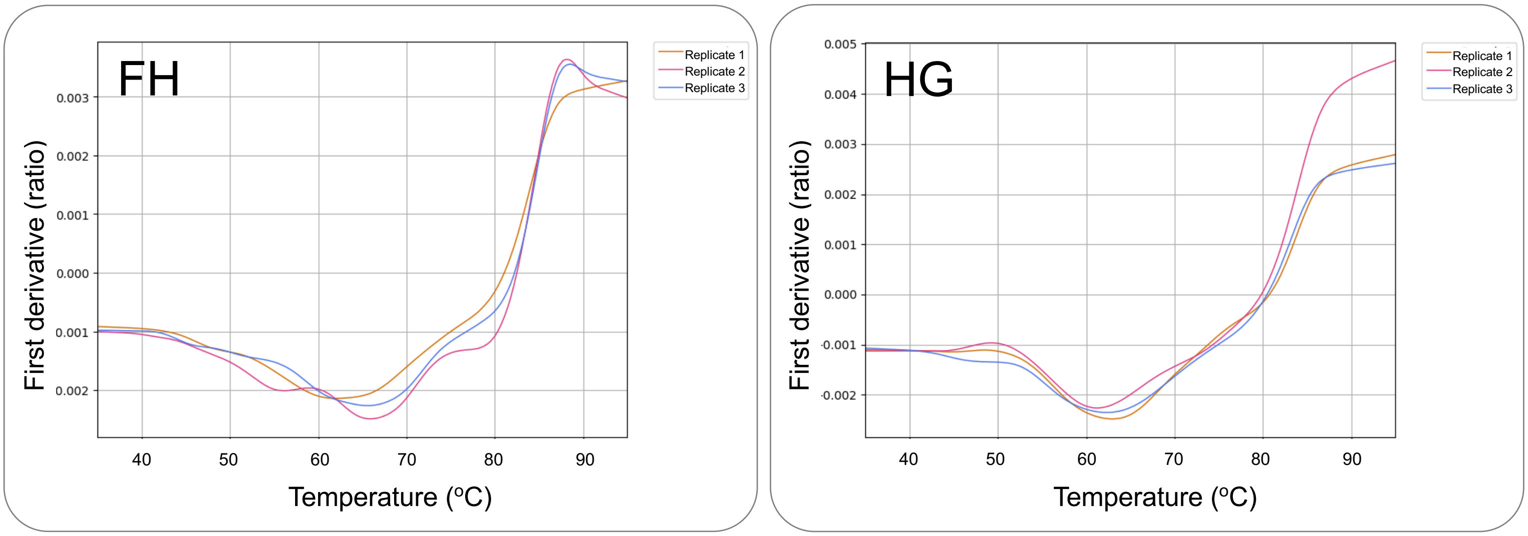

Figure S3. Simulation of SOD1 trimer (yellow; model from Proctor et al., 2016) and septin-7 (red; PDB ID 6N0B) from ClusPro model docking through discrete molecular dynamics. Protein binding simulation overlayed with the septin-2G/Septin-6/septin-7 hetero-hexamer (PDB ID 7M6J). (orange, green, blue).

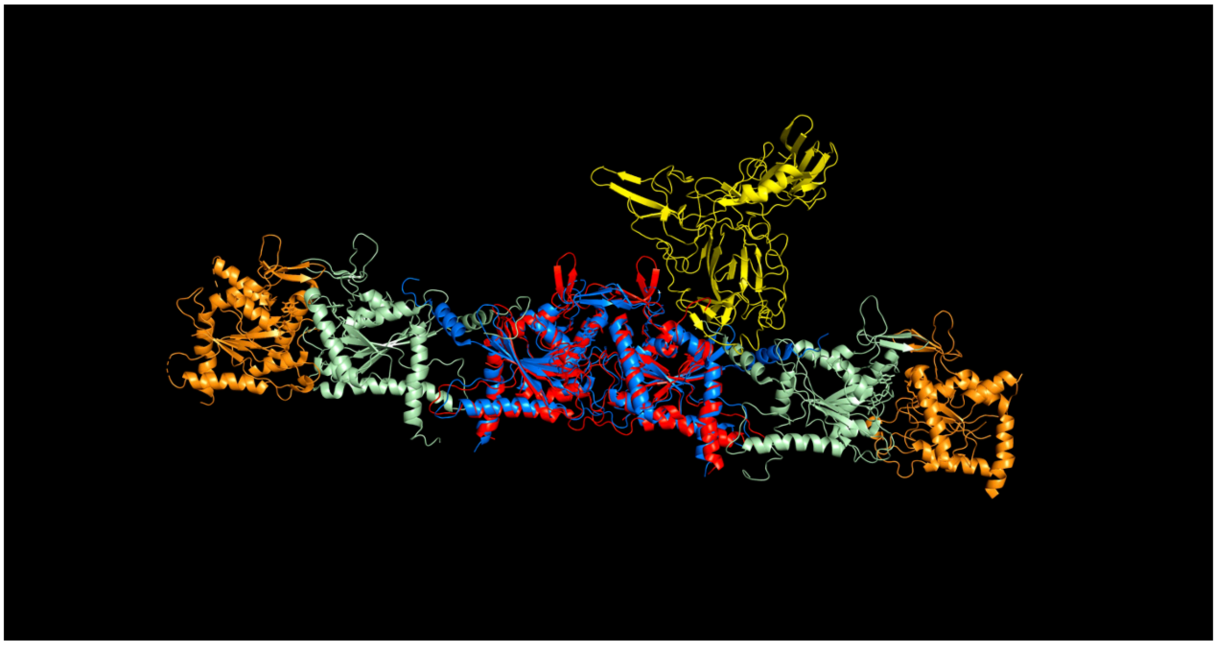

(video file separate)

Figure S4. Estimation of SOD1 trimer concentration in the brain and physiological relevance to MST $K_d$ measurements. 1. We used the PAXdb 5.0 to obtain concentrations of septin-7 and SOD1 trimer in the brain. Septin-7 concentration in the brain is 809 ppm which is about 16 uM. SOD1 concentration in the brain is about 3488 ppm which is about 218 uM. 2. From Khare et al. in 2004 observed about 12% aggregation of SOD1 in non-dialysis conditions at SOD1 [30 uM]. Hnath and Dokholyan in 2022 determined that with ALS mutation A4V, there is about 50/50 trimer/aggregates. So, we estimate that 6% of SOD1 is trimer and that in the brain about 13 uM could be trimers. 3. We estimated the amount of septin-7 bound by trimer using the [bound] = [trimer] [SEPT7] / $K_d$ + [SEPT7]. 4. We interpret the $K_d$ measurements (FH median $K_d$ of 7.48from SOD1 trimer and septin-7 MST experiments to be physiologically relevant since about 8.8 uM of the total 16 uM of septin-7 in the brain may be occupied by SOD1 trimers. These are conservative estimates. Image is created with BioRender.com.

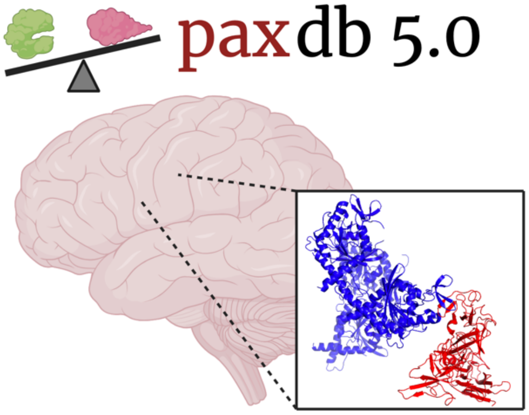


1. PAXdb 5.0 concentration estimation in the brain
[SEPT7] 16 uM
[SOD1] 218 uM
2. Estimate SOD1 trimer concentration aggregation percent calculation from Khare et al. (2004) and trimer:aggregates determination from Hnath and Dokholyan (2022)
    in SOD1 [30 uM] there is about 12% aggregation
    50/50 trimer:aggregates
    6% trimer
SOD1 brain [218 uM] * 6% as trimers = SOD1 trimers [13 uM]
3. [bound] = [trimer] [SEPT7] / $K_d$ + [SEPT7]
    [bound] = [13 uM] [16 uM] / 7.48 + [16 uM]
    [bound] = 8.8 uM
4. interpretation: about 8.8 uM out of the total 16 uM of SEPT7 in the brain may be occupied by SOD1 trimers.

Figure S5. Confocal images staining stably expressing SOD1 trimer under doxycycline control in NSC-34s. We stained the stable NSC-34 cells with C4F6 in red which selectively stains misfolded SOD1 in WT and trimer expressing cells. Septin-7 is stained in green. We also stained a set of NSC-34 cells with WT SOD1 antibody in WT and trimer expressing cells. We stained the nuclei with Hoechst (blue). We performed co-localization analysis of WT of misfolded SOD1 with septin-7.

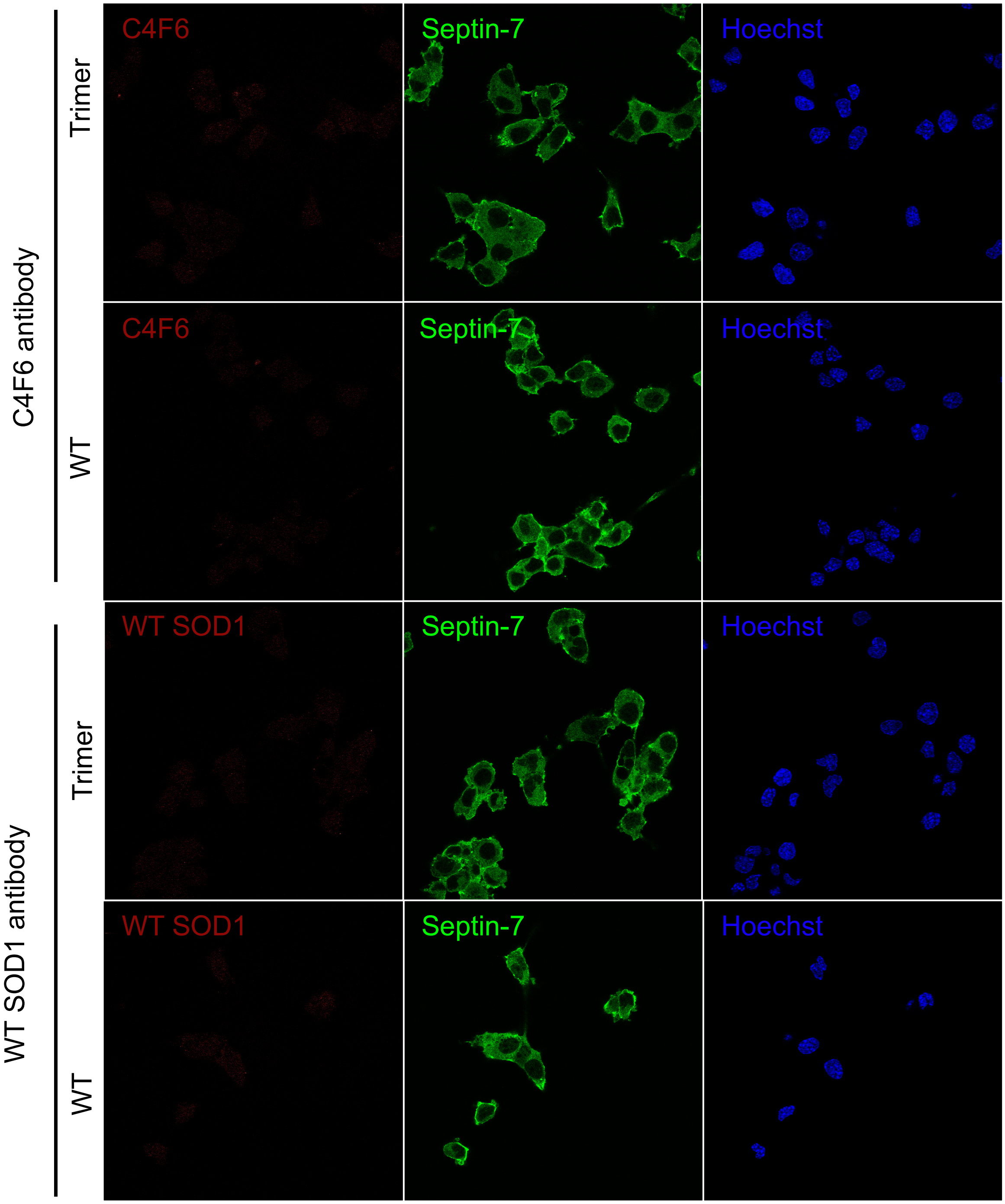
